# Striatal dopamine regulates sleep states and narcolepsy-cataplexy

**DOI:** 10.1101/2023.05.30.542872

**Authors:** Brandon A. Toth, Katie S. Chang, Christian R. Burgess

## Abstract

Disruptions to sleep can be debilitating and have a severe effect on daily life. Patients with the sleep disorder narcolepsy suffer from excessive daytime sleepiness, disrupted nighttime sleep, and cataplexy - the abrupt loss of postural muscle tone (atonia) during wakefulness, often triggered by strong emotion. The dopamine (DA) system is implicated in both sleep-wake states and cataplexy, but little is known about the function of DA release in the striatum - a major output region of midbrain DA neurons - and sleep disorders. To better characterize the function and pattern of DA release in sleepiness and cataplexy, we combined optogenetics, fiber photometry, and sleep recordings in a murine model of narcolepsy (orexin^−/−^; OX KO) and in wildtype mice. Recording DA release in the ventral striatum revealed OX-independent changes across sleep-wake states as well as striking increases in DA release in the ventral, but not dorsal, striatum prior to cataplexy onset. Tonic low frequency stimulation of ventral tegmental efferents in the ventral striatum suppressed both cataplexy and REM sleep, while phasic high frequency stimulation increased cataplexy propensity and decreased the latency to rapid eye movement (REM) sleep. Together, our findings demonstrate a functional role of DA release in the striatum in regulating cataplexy and REM sleep.

## Introduction

Narcolepsy is a prevalent and debilitating rapid eye movement (REM) sleep-related disorder, characterized by excessive daytime sleepiness (EDS) and disrupted nighttime sleep (DNS), in addition to cataplexy - the inappropriate recruitment of brainstem REM sleep muscle atonia-generating circuitry in response to positively-valenced stimuli^1–5^. Narcolepsy type 1 is caused by the loss of lateral hypothalamic neurons expressing orexin/hypocretin (OX), a peptide heavily implicated in the maintenance of wakefulness^6,7^. Orexin mediates wake-promoting effects in part through coordinating the release of monoamines, including dopamine (DA)^5,8,9^. Midbrain DA neurons in the ventral tegmental area (VTA) and substantia nigra pars compacta (SNc) have been implicated in a variety of motivated behaviors and neurological disorders^10–13^ and have also recently been shown to drive arousal states, in addition to having differing activity patterns across sleep-wake states^14–20^. Collectively, this suggests that aberrant DA release could underlie disorders of sleep and arousal.

One of the key targets of midbrain DA neurons are inhibitory medium spiny neurons in the striatum, with which these neurons have reciprocal connectivity^21–26^. The striatum is composed of functionally and anatomically distinct regions^27^, and the activity of DA terminals differs between dorsal and ventral striatal structures^28^. SNc projections to the dorsolateral striatum (DLS) are generally associated with motor control and habit formation, while VTA projections to ventral regions such as the nucleus accumbens (NAc) and its core (NAcc) and shell (NAcSh) subregions are associated with reward and learning^28–31^. During REM sleep, activity in the ventral striatum increases to levels similar to wakefulness, with DA release also increasing to wake-like levels^18,32,33^. Furthermore, relative to healthy subjects, patients with narcolepsy show increased activity in the ventral striatum in response to cataplexy-evoking stimuli^34^, while lesions to this region attenuate cataplexy frequency^35^. Together, these data indicate that the striatum may play a role in the regulation of REM sleep and cataplexy. However, the functional role of DA release in the striatum with respect to EDS, DNS, and cataplexy, in addition to the relationship between the OX system and DA release remains unclear.

In the present study, we used optogenetics and fiber photometry to examine the pattern and function of DA release in different regions of the striatum during changes in sleep-wake states and cataplexy in a murine model of narcolepsy (orexin^−/−^; OX KO). First, we show that while there are clear differences in DA tone across sleep-wake states, there is no difference in DA release within sleep-wake states between OX KO and wildtype (WT) mice. Second, we demonstrate that there is a striking increase in DA release prior to cataplexy onset in both the NAcc and NAcSh, but not in the DLS. Finally, we found that low frequency, tonic stimulation of midbrain efferents in both the NAcc and NAcSh reduced cataplexy and REM sleep occurrence. Conversely, high frequency, phasic stimulation of VTA efferents in the NAcc, but not NAcSh, decreased the latency to REM sleep and increased cataplexy propensity. Together, these data indicate that DA release in the striatum across sleep-wake states is OX-independent. Furthermore, we conclude that DA release in the ventral striatum has a functional role in cataplexy and REM sleep.

## Results

### NAcc DA dynamics do not differ between WT and OX KO mice

In both human and murine narcolepsy, the most commonly observed phenotype is behavioral state instability, manifesting as low thresholds to transitions between sleep-wake states^4,5,36^. To accurately assess DA release dynamics across sleep-wake states, dLight1.1^37^ was expressed in the NAcc of WT and OX KO mice, and an optic fiber was implanted to permit *in vivo* fiber photometry recordings of DA tone (Figure 1A, B; Figure S1A). Following recovery from surgery, mice were habituated to recording chambers which featured electroencephalography/electromyography (EEG/EMG) recording tethers, a fiber optic cable for photometry, and in-cage running wheel (Figure S1B).

**Figure 1.**
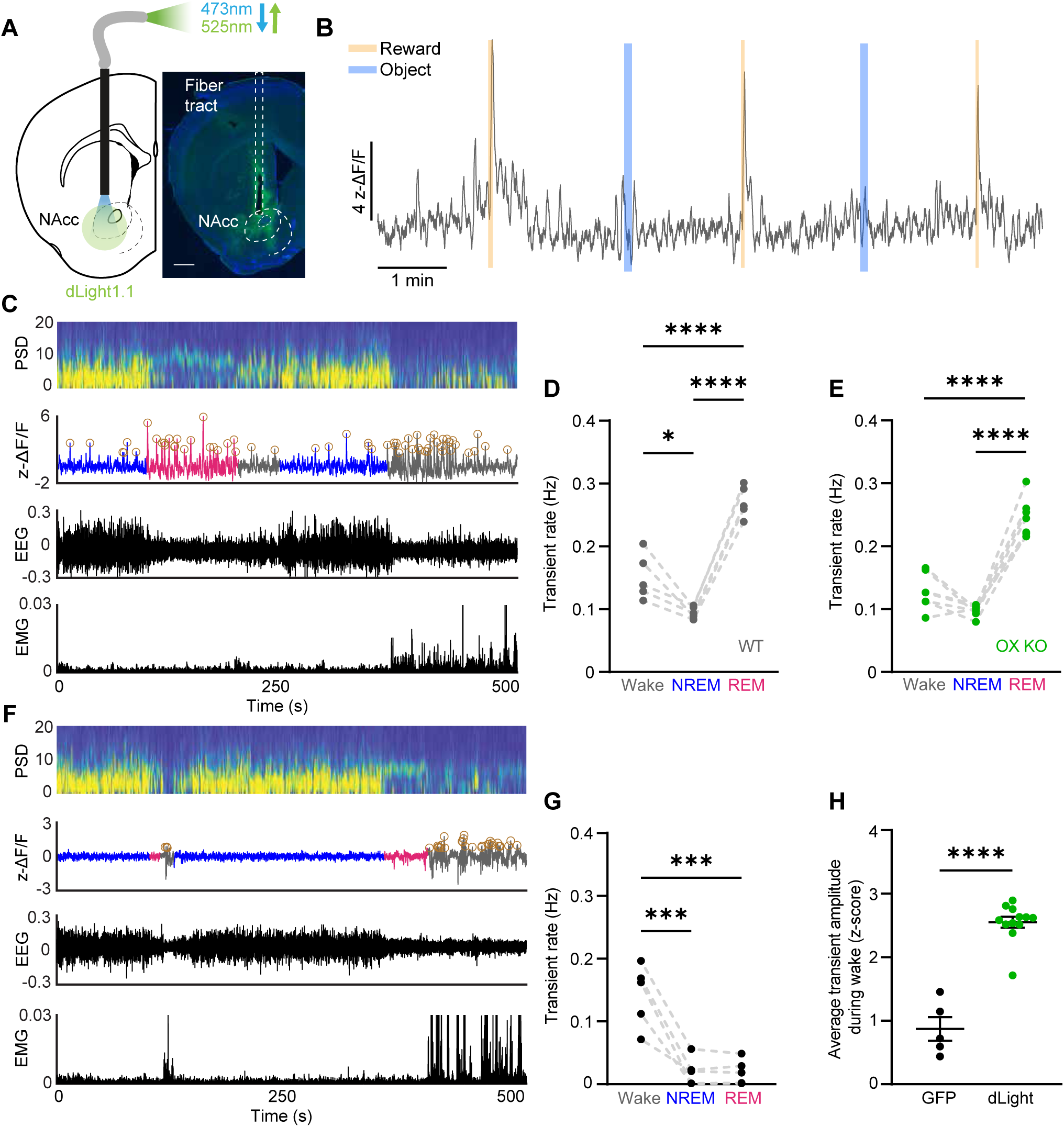
DA release in the NAcc fluctuates across sleep-wake states independent of OX expression. (A) Schematic of photometry recording in the NAcc (left). Expression of dLight in the NAcc (green: dLight, blue: DAPI). Scale bar equal to 400 µm (right). (B) ΔF/F trace during a recording session for a representative WT animal. Reward and object presentations were alternated throughout the session. DA release was elevated in response to reward, but not to a non-food object. (C) Representative power spectrogram, ΔF/F trace, EEG, and EMG across sleep-wake states in an OX KO animal. Circles around peaks represent DA transients. (D - E) Transient rate during Wake, NREM, and REM sleep in WT (D; n = 5) and OX KO (E; n = 7) mice. (F) Representative power spectrogram, ΔF/F trace, EEG, and EMG across sleep-wake states in a GFP-expressing animal. Circles around peaks represent automatically detected motion artifact induced transients in the GFP (n = 5) signal. (G) Transient rate during wake, NREM, and REM sleep in GFP-expressing mice. (H) Average fluorescence of detected transient peaks during wakefulness. While transients are detected in the GFP signal, they are significantly smaller than real DA transients detected in dLight recordings. All data represented as the mean ± SEM. *p<0.05, ***p<0.001, ****p<0.0001; one-way ANOVA with Tukey post-hoc comparison test (D, E, G), unpaired t-test (H).

We first confirmed previous behavioral characterizations of OX KO mice. While total time spent within sleep states was similar between OX KO (n = 8) and WT (n = 5) mice, OX KO mice exhibited shorter bout durations, increased numbers of bouts of each state, and increased brief arousals from non-REM (NREM) sleep (Figure S1C-F). We next evaluated changes in DA release across sleep-wake state transitions during a 12 h light period. Transitions from Wake-to-NREM showed little change, while transitions from NREM-to-Wake elicited a robust increase in fluorescence (Figure S1G, H, K, L). However, in both mice expressing dLight and GFP (n =5), we also observed a decrease in the fluorescent signal during NREM-to-REM transitions and subsequent rebound during REM-to-Wake transitions (Figure S1I, J, M, N). As this likely reflects a form of fluorescent artifact^38,39^, slow changes in the photometry signal were removed and we analyzed the rate of DA transients as a more faithful reflection of DA release within states, particularly REM sleep (see Methods for a detailed discussion).

We then asked if the loss of OX expression altered NAcc DA transient rates within sleep-wake states, as OX neurons directly affect the activity of VTA DA neurons^8,9^. To our surprise however, there were no significant differences in transient rates between WT and OX KO mice in any state. In NREM sleep, DA release was lowest in both WT and OX KO mice, relative to wakefulness and REM sleep (Figure 1C-E). DA transient rates were most pronounced during REM sleep, consistent with prior literature suggesting that DA release in the NAcc increases during REM sleep^15,18^. We also detected ‘transients’ in control mice expressing GFP during wakefulness, though this is likely due to motion artifacts, as 1) unlike dLight, GFP signals showed no transient activity during NREM and REM sleep and 2) detected transient-like events were significantly smaller in amplitude than those detected in dLight-expressing mice (Figure 1F-H). Taken together, these findings suggest that while there are profound behavioral differences in sleep states between WT and OX KO mice, DA release in the NAcc does not underlie behavioral state instability in narcolepsy. Furthermore, they demonstrate that DA release in the NAcc may have a functional role in the regulation of REM sleep.

### DA release in the NAcc increases prior to cataplexy onset

Cataplexy is often described as the intrusion of REM sleep muscle atonia into wakefulness, often in response to positively valenced stimuli^4,5^. As DA release in the NAcc is associated with positive valence states, we hypothesized that DA release would increase prior to the onset of cataplexy. To increase the number of cataplexy episodes, recordings were done during 12 h dark periods with chocolate availability, as chocolate has been shown to increase cataplexy propensity^2,3,35^ (Figure S2B). Consistent with previous reports^36^, episodes of cataplexy had increased theta rhythmicity and there was no difference in the spectral power between REM sleep and cataplexy (Figure 2A, B). Furthermore, there was no difference when comparing DA transient rates in REM sleep and cataplexy (n = 7; Figure 2C). Consistent with our hypothesis, prior to the onset of cataplexy we observed a striking increase in DA release, returning to baseline following transitions back into wakefulness (n = 7; Figure 2D-F, Figure S2A). These dynamics were similar whether recordings were done in the presence or absence of chocolate (Figure S2C, D). These results demonstrate that the NAcc may be a key functional node that DA acts on to regulate cataplexy. Notably, the increase in DA release preceded the change in state, demonstrated by the rise in DA preceding the change in theta rhythmicity, suggesting that DA release in the NAcc may drive transitions into cataplexy.

**Figure 2.**
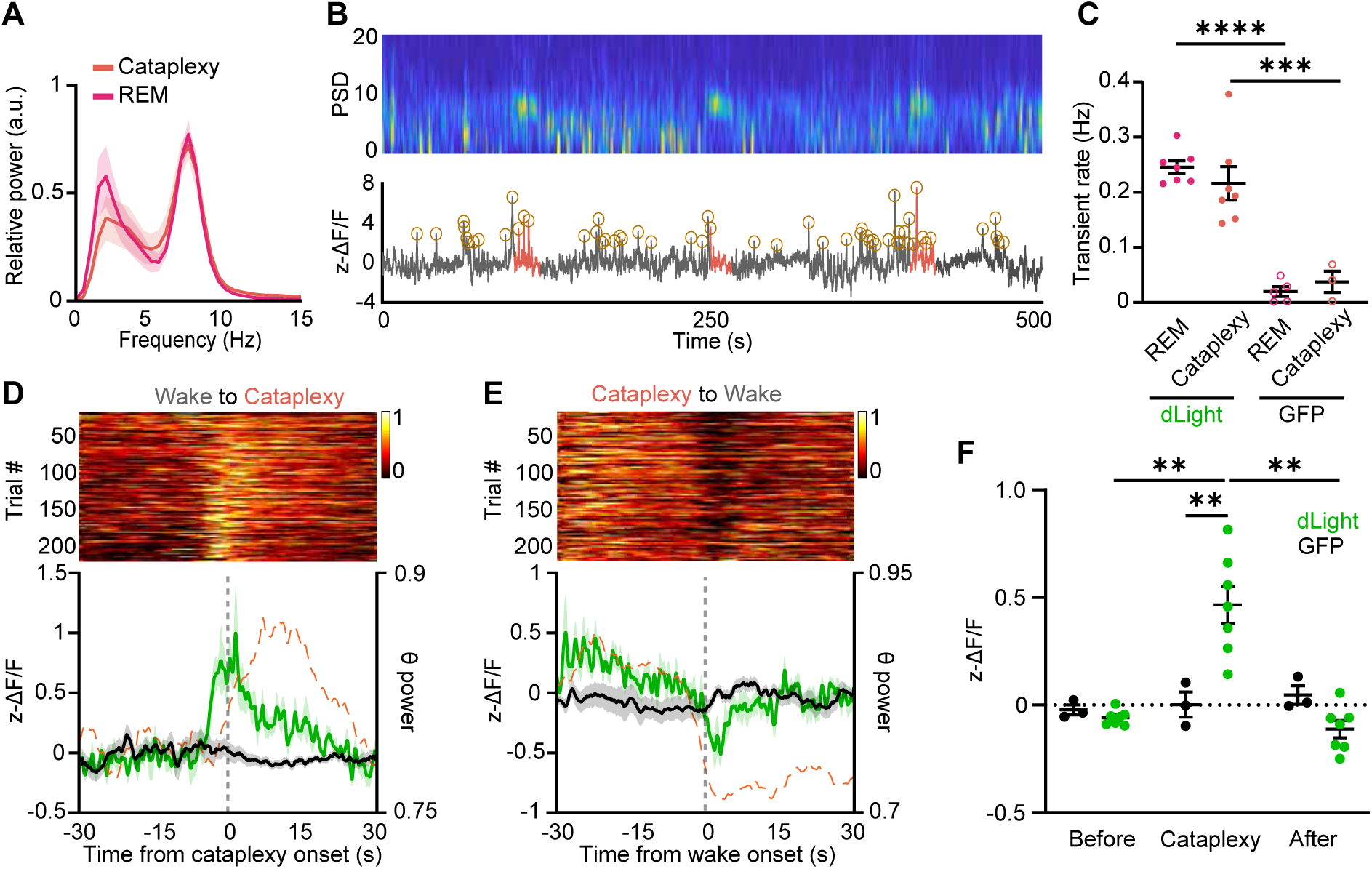
DA release in the NAcc is elevated prior to cataplexy onset. (A) Power spectral density of the EEG during REM sleep and cataplexy. (B) Representative power spectrogram and ΔF/F trace during several transitions into cataplexy in an OX KO animal. Circles around peaks represent automatically detected DA transients. (C) Transient rate during REM sleep and cataplexy in dLight- and GFP-expressing mice (dLight, n = 6; GFP, n = 3-5). (D) Single-trial cataplexy-onset-triggered time courses of DA release in the NAcc, sorted by response amplitude at the onset of cataplexy (top). Average dLight fluorescence (solid green trace) and normalized theta power (dashed orange trace) aligned Wake-to-Cataplexy transitions (bottom). (E) Same as (D) but for Cataplexy-to-Wake transitions. (F) Average dLight fluorescence in the period prior to, during, and after cataplexy (n = 7). All data represented as the mean ± SEM. **p<0.01; two-way ANOVA with Sidac post-hoc comparison test (C, F).

### DA release in the striatum differs along the dorsal-ventral axis during sleep-wake transitions and cataplexy

The striatum is composed of functionally and anatomically distinct regions, where DA release drives different behaviors along the dorsal-ventral axis^27^. DA release in the dorsal striatum has been linked to locomotion, while DA release in the ventral striatum encodes information about reward^28^. In light of these distinctions, to further characterize DA release in the striatum during sleep-wake states and cataplexy, we next expressed dLight1.1 in the DLS (n = 5) or NAcSh (n = 5) of OX KO mice and recorded DA release throughout the 12 h light period (Figure 3A-C). Similar to the NAcc, we observed a fluorescent decrease during REM sleep in both the DLS and NAcSh (Figure S3). As such, we evaluated DA dynamics across states by looking at DA transients.

**Figure 3.**
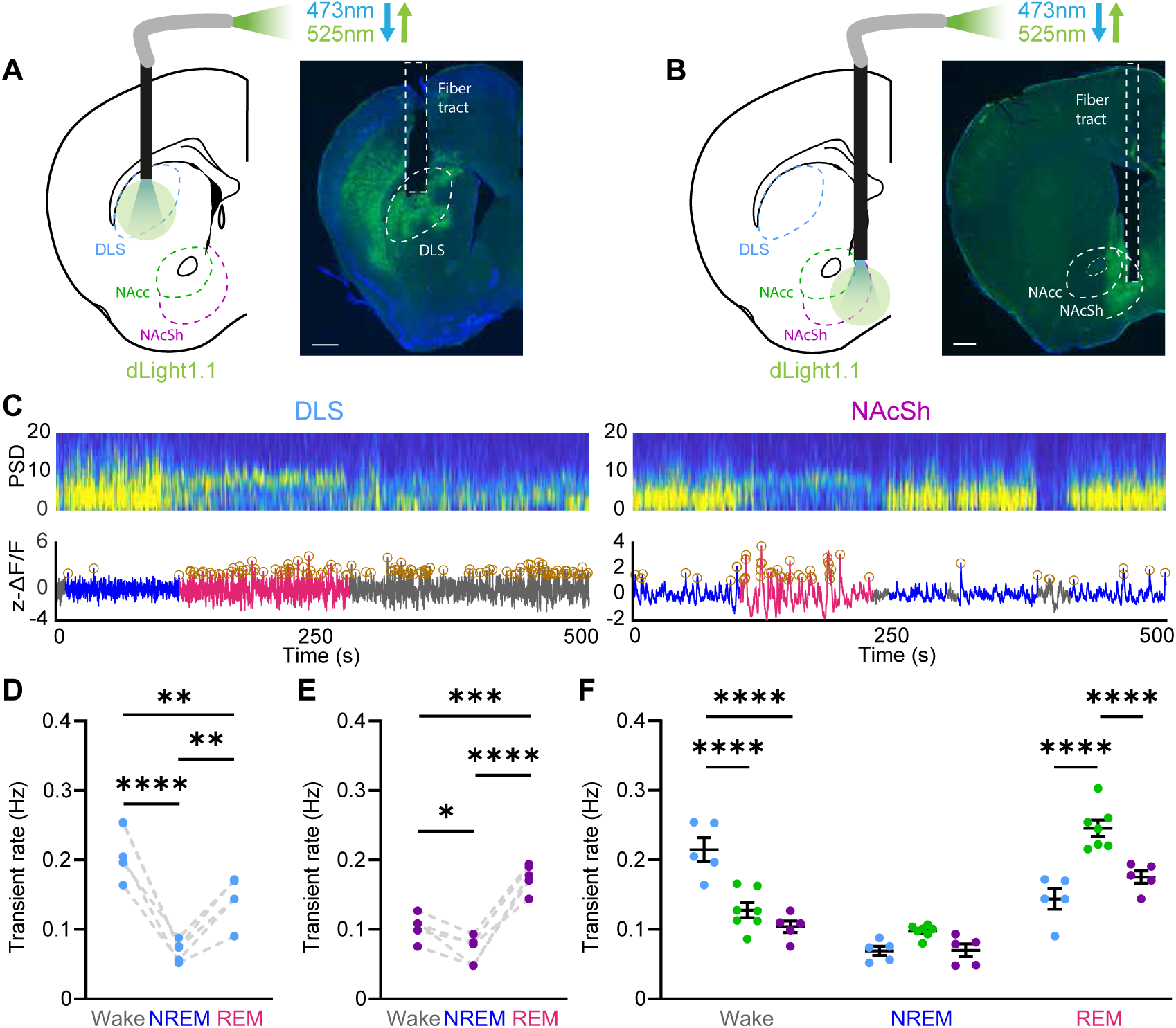
DA release differs across striatal regions within sleep-wake states. (A, B) Schematic and expression of dLight in the DLS (A) and NAcSh (B) (green: dLight, blue: DAPI) Scale bars equal to 400 µm. (C) Representative power spectrogram and ΔF/F trace in the DLS (left) and NAcSh (right). (D, E) DA transient rate across sleep-wake states in the DLS (D) and NAcSh (E). (F) DA transient rate during sleep-wake states across different striatal regions (n = 5 for NAcSh and DLS, n = 7 for NAcc). All data represented as the mean ± SEM. *p<0.05, **p<0.01, ***p<0.001, ****p<0.0001; one-way ANOVA with Tukey post-hoc comparison test (D, E), two-way ANOVA with Sidak post-hoc comparison test (F).

Consistent with the proposed role for the DLS in locomotion, we found that DA transient rates were highest in wake, relative to both NREM and REM sleep (Figure 3D). Interestingly, DA release was also elevated during REM sleep relative to NREM sleep. In the NAcSh, transients were highest in REM sleep, relative to both wake and NREM sleep (Figure 3E). When evaluating within state DA release across regions, during wakefulness, transients in the DLS were significantly elevated relative to both the NAcc and NAcSh, while during REM sleep, transient rate in the NAcc was significantly higher than in the DLS and NAcSh (Figure 3F). Thus, DA release in the dorsal striatum is more biased towards wakefulness, whereas DA release in the ventral striatum, particularly in the NAcc, is biased towards REM sleep.

Although transients in the DLS were elevated during wakefulness relative to the NAcc and NAcSh, across NREM-to-Wake transitions, there were no significant differences between striatal regions, with all regions having increased release across the transition (Figure S3E). This however was not the case for brief arousals from NREM sleep - DA release in the DLS was elevated during brief arousals, whereas it was suppressed in the NAcc and NAcSh (Figure S3B, C). This suggests that with regard to wakefulness, DA release in the DLS may reflect general arousal/locomotion, while DA release is only elevated in the ventral striatum during sustained arousal and may reflect the content of wakefulness.

We next asked if DA release during episodes of cataplexy in the DLS and NAcSh followed the same patterns observed in the NAcc. Contrary to the robust dynamics seen at the onset of cataplexy in the NAcc, little change was observed in the DLS (Figure 4A-D, Figure S4A,B). However, following transitions back into wakefulness, rather than a suppression of DA as seen in the NAcc, DA release increased above baseline, perhaps reflecting the role for DA release in the DLS in locomotion. In the NAcSh, similar to the NAcc, we saw a robust increase in DA release prior to the onset of cataplexy (Figure 4E-H, Figure S4C,D). Furthermore, in the wakefulness following cataplexy, relative to the baseline period, DA release in the NAcSh was significantly suppressed (Figure 4G, H). When evaluating DA release across striatal regions during cataplexy, while DA transient rates in the NAcc and NAcSh were significantly elevated relative to the DLS, overall DA release increases along the dorsal-ventral axis (Figure 4I-L, Figure S4E). Together, these results illustrate a dorsal-ventral pattern of release, and perhaps differing roles, for DA in the striatum during sleep-wake states and suggests that DA release in the NAcc and NAcSh may engage circuitry to promote cataplexy.

**Figure 4.**
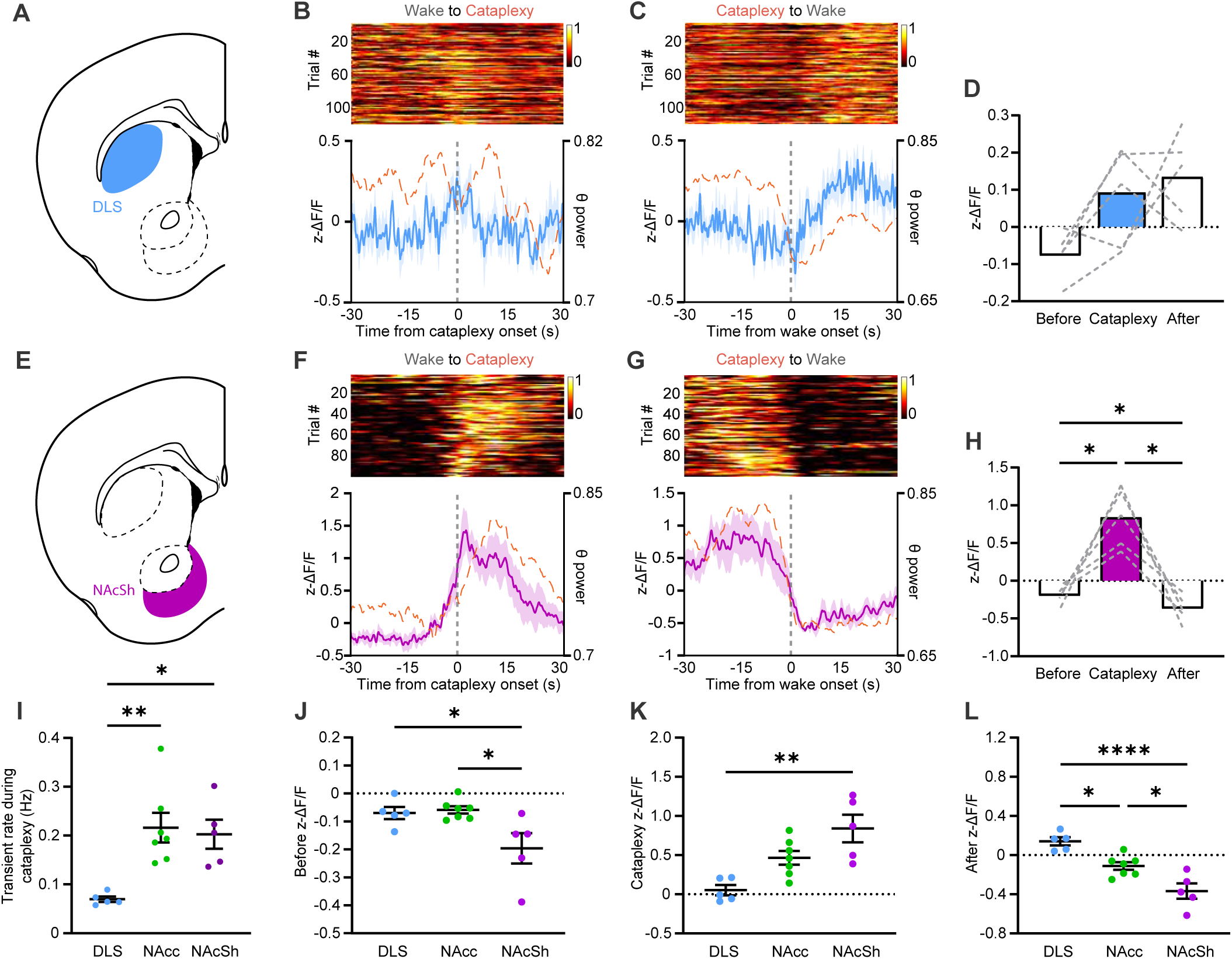
DA release at cataplexy onset is more pronounced in the ventral striatum. (A) Schematic representation of the DLS. (B) Single-trial cataplexy-onset-triggered time courses of DA release in the DLS (n = 5), sorted by response amplitude at onset of cataplexy (top). dLight fluorescence (solid blue trace) and normalized theta power (dashed orange trace) aligned to Wake-to-Cataplexy transitions (bottom). (C) Same as (B) but for Cataplexy-to-Wake transitions. (D) Average fluorescence in the period prior to, during, or after cataplexy. (E) Schematic representation of the NAcSh. (F - H) Same as (B - D) but for DA release in the NAcSh (n = 5) during cataplexy. (I) DA transient rate across striatal regions during cataplexy. (J - L) dLight fluorescence before (J), during (K), and after (L) cataplexy episodes in the DLS, NAcc, and NAcSh. All data represented as the mean ± SEM. *p<0.05, **p<0.01, ****p<0.0001; RM one-way ANOVA with Tukey post-hoc comparison test (D, H, I), one-way ANOVA with Tukey post-hoc comparison test (J - L).

### DA release during cataplexy occurs independent of the preceding behavior

In both human narcolepsy patients and murine models of narcolepsy, a number of behaviors and experiences can elicit cataplexy^40,41^. Therefore, we asked if changes in DA release during cataplexy occurred due to the behavior the animal was engaged in prior to the onset of cataplexy. Using videography, we scored 6 independent behaviors that occurred immediately before an episode of cataplexy: burrowing, feeding, licking, grooming, running wheel activity, and spontaneous (the lack of a definable behavior). Across all mice used in recordings in the DLS, NAcc, and NAcSh, consistent with previous characterizations in OX KO mice^40^, approximately 90% of cataplexy episodes were preceded by grooming, running, or spontaneous activity (Figure 5A). We then assessed DA release dynamics during cataplexy preceded by these more prominent behaviors. In the DLS, as expected, there was no significant change at cataplexy onset for any behavior (Figure 5B). In both the NAcc and NAcSh however, cataplexy preceded by any behavior had similar DA release dynamics, with significantly increased DA release during cataplexy relative to baseline (Figure 5C, D). These findings indicate that DA release in the NAcc and NAcSh is a defining feature of all cataplexy episodes, regardless of prior behavior.

**Figure 5.**
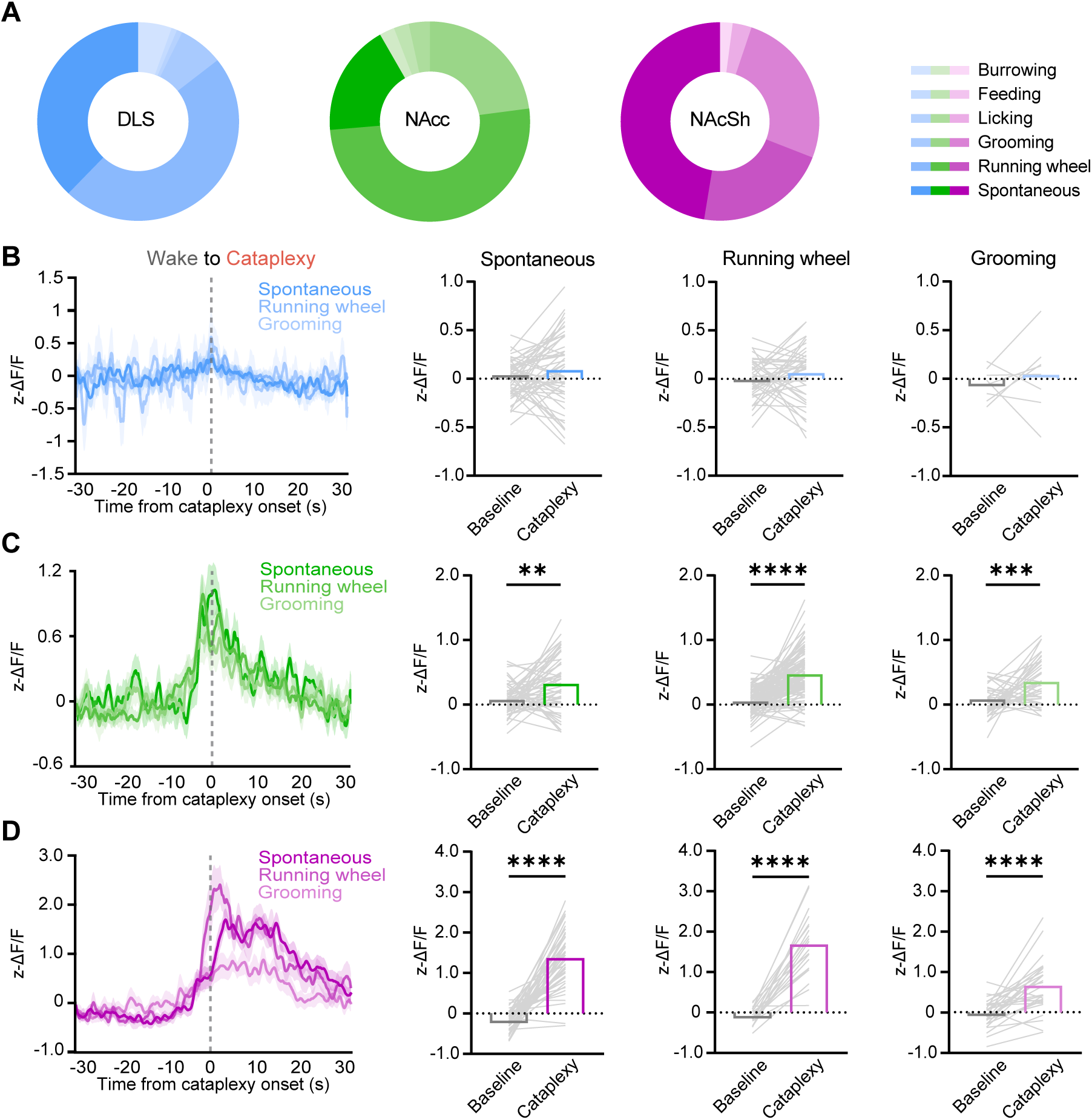
DA release at cataplexy onset is independent of prior behavior. (A) Pie charts of the different behaviors observed to precede episodes of cataplexy in DLS (n = 111), NAcc (n = 205), and NAcSh (n = 96) dLight-expressing OX KO mice. (B) Overlay of dLight fluorescence in the DLS aligned to Wake-to-Cataplexy transitions, separated by different behaviors that preceded cataplexy. Average dLight fluorescence prior to and during cataplexy preceded by spontaneous, running wheel, and grooming activity. (C, D) Same as (B) but for DA release in the (C) NAcc and (D) NAcSh during cataplexy. All data represented as the mean ± SEM. *p<0.05, **p<0.01, ****p<0.0001; paired t-test (C, D).

### Optogenetic stimulation of VTA efferents in the ventral striatum controls REM sleep and cataplexy

Given the striking DA release seen prior to cataplexy onset in the ventral striatum, we anticipated that *in vivo* manipulation of the VTA→NAcc/NAcSh circuit in OX KO mice would alter cataplexy frequency. VTA DA neurons generally exhibit two different firing states: low-frequency tonic activity and high-frequency phasic activity^42^. Paradoxically, sustained DA release in the ventral striatum results in inhibition of phasic DA activity^43–48^, suggesting that tonic vs phasic stimulation may impact cataplexy propensity differently. To specifically target NAcc/NAcSh-projecting neurons in the VTA, we unilaterally injected a retrograde AAV expressing Cre recombinase into either the NAcc (n = 5-6) or NAcSh (n = 5-6) and injected a Cre-dependent AAV expressing ChrimsonR in the VTA (Figure 6A-C).

**Figure 6.**
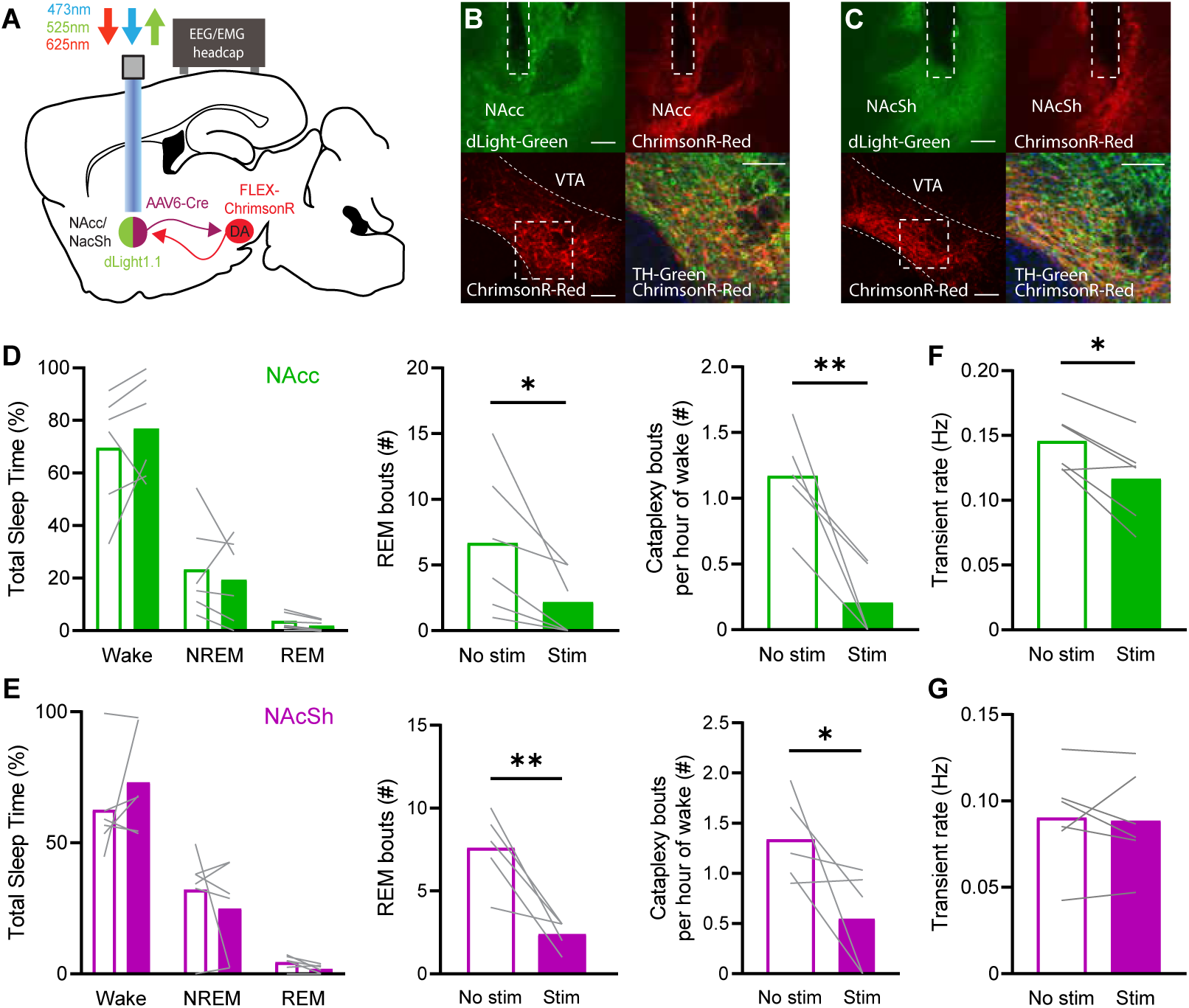
Tonic VTA efferent stimulation in the NAcc and NAcSh suppresses REM sleep and cataplexy. (A) Schematic of circuit-specific optogenetic stimulation. (B) Representative histology of dLight and ChrimsonR terminals expressed in the NAcc (top) and ChrimsonR/tyrosine hydroxylase (TH) colocalization in the VTA (bottom). Scale bars are equal to 100 µm. (C) Same as (B), but for the NAcSh. (D) Effect of 2 h continuous 2 Hz stimulation in the NAcc (n = 5-6) on total sleep time, number of REM bouts, and number of cataplexy bouts per hour of time spent awake. (E) Same as (D) but for continuous 2 Hz stimulation in the NAcSh (n = 5-6). (F, G) Wake transient rate during tonic stimulation in the (F) NAcc and (G) NAcSh. All data represented as the mean ± SEM. *p<0.05, **p<0.01; paired t-test (D, E, F, G).

We first performed continuous 2 Hz stimulation of VTA efferents in the NAcc and NAcSh of OX KO mice over a 2 h recording session and found that, relative to recordings with no stimulation, while there was no effect on total sleep time, bouts of REM sleep and cataplexy were strongly suppressed (Figure 6D, E). Notably, we found that transient DA release during waking in the NAcc, but not NAcSh, was indeed inhibited during tonic stimulation (Figure 6F, G). Thus, tonic DA release in the ventral striatum has a suppressive effect on REM sleep and cataplexy, potentially by altering phasic release.

Since the increase in DA release in both the NAcc and NAcSh preceded entrances into cataplexy (Figure 2D, 4F), we hypothesized that driving phasic DA release in the NAcc and NAcSh would increase cataplexy frequency. Furthermore, as we observed increased DA release during REM sleep (Figure 1E, 3E), we anticipated that phasic DA release during NREM would also increase transitions into REM sleep. However, when stimulating efferents using a brief, 20 Hz stimulation protocol (5 ms pulses at 20 Hz for 5 s, with a variable 8-10 min ITI), despite increased DA release, there was little effect on sleep-wake architecture, cataplexy, or latency to wake, REM, or cataplexy with either VTA→NAcc or VTA→NAcSh terminal stimulation (Figure S5). Therefore, we concluded that brief phasic DA release in the ventral striatum is not sufficient to induce changes in sleep-wake states and cataplexy.

To better recreate the endogenous DA release dynamics observed during cataplexy, we developed a variable stimulation protocol. In this protocol, stimulation began at 10 Hz and gradually increased to 40 Hz over the course of 20 s, with a variable 8-10 min ITI (Figure 7A). Stimulation elicited a robust increase in DA release in both the NAcc and NAcSh that closely resembled the time course of release during cataplexy (Figure 7B). When stimulating VTA→NAcc terminals, while there was no change in the overall sleep-wake architecture, we saw a significant increase in cataplexy propensity, with no change in the latency to cataplexy onset (Figure 7C, D). When stimulation occurred during NREM sleep, we found that the latency to both wake and REM sleep decreased. Interestingly, despite a robust increase in DA release, we saw no significant increase in cataplexy when stimulating VTA→NAcSh terminals (Figure 7E). Similarly, there was no effect on sleep state architecture or latencies (Figure 7F). Taken together, tonic vs phasic DA release in the NAcc differentially regulates cataplexy propensity and alters sleep-wake architecture.

**Figure 7.**
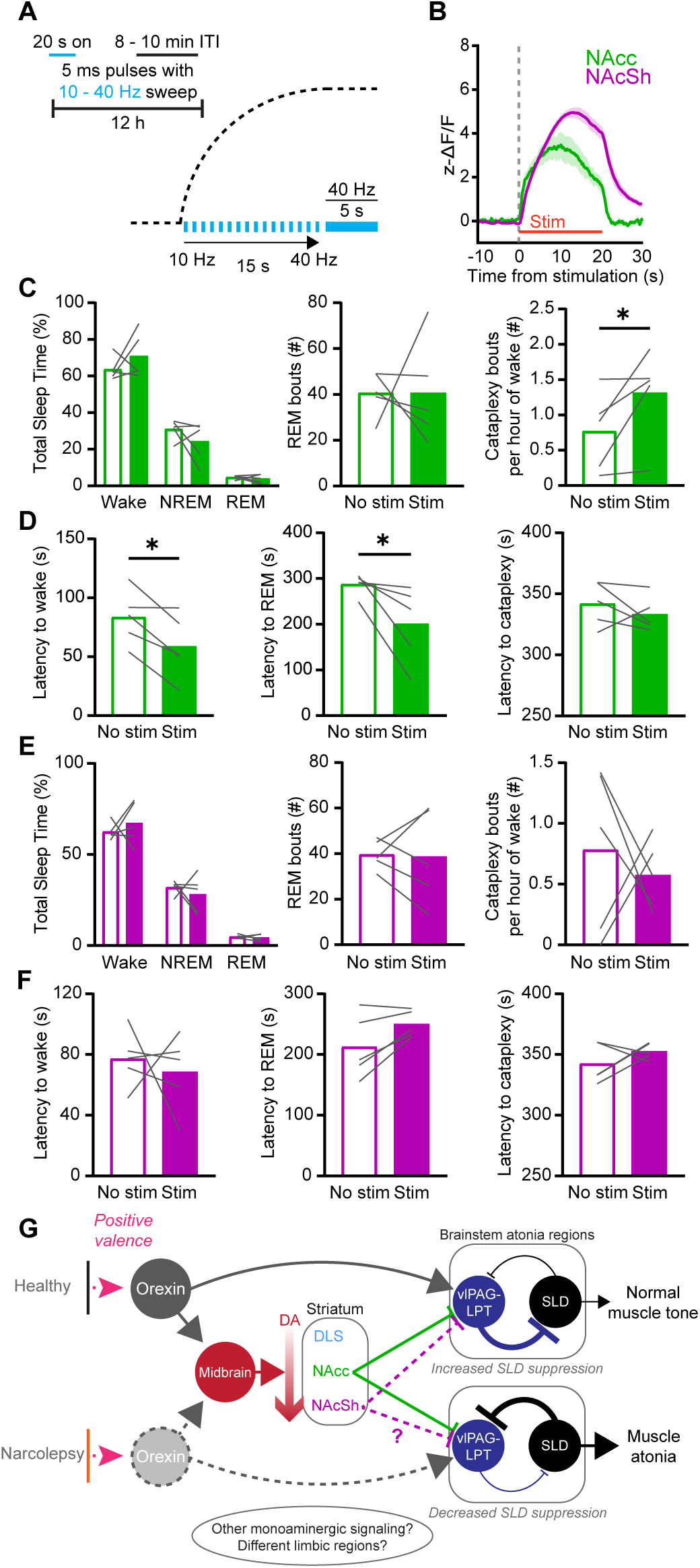
Prolonged phasic stimulation of VTA efferents in the NAcc promotes cataplexy. (A) Schematic of the 12 h variable sweep stimulation paradigm. (B)dLight response to variable sweep stimulation in the NAcc and NAcSh (n = 5 in both groups). (C) Effect of variable sweep stimulation in the NAcc on total sleep time, number of REM bouts, and number of cataplexy bouts per hour of time spent awake. (D) Effect of variable sweep stimulation in the NAcc on latency to Wake, REM, and Cataplexy. (E, F) Same as (C, D), but for the NAcSh. (G) Proposed circuit for limbic regulation of cataplexy/muscle atonia in narcolepsy. Optogenetically evoked phasic DA release in the NAcc increased cataplexy propensity but was not sufficient to directly trigger immediate entrances into cataplexy, suggesting that other neuromodulators and brain regions, such as the NAcSh, may act in concert with DA circuits. All data represented as the mean ± SEM. *p<0.05; paired t-test.

## Discussion

Loss of the OX system introduces profound effects on sleep-wake architecture. Yet while regions of the striatum and the DA system are both known to regulate fundamental behaviors and arousal states, few studies have demonstrated, with high temporal and spatial resolution, how the two may contribute to disorders of sleep and arousal, such as narcolepsy. Combining finescale recording and manipulation of DA release, we have characterized changes in DA release in the striatum across sleep-wake states that are region-specific and OX-independent. Furthermore, we not only characterized the endogenous release profile of DA in the striatum during REM sleep and cataplexy, but also describe a functional role for DA release in the ventral striatum in modulating the frequency of both states. These findings offer new insights into the relationship between OX and DA release in sleep states and illustrate the involvement of the striatal DA in regulating cataplexy.

### Striatal DA release differs across sleep states, independent of OX signaling

Both VTA and SNc DA neurons have previously been shown to have differing activity patterns during sleep-wake states^16,20,49^. Yet while OX has long been considered to be integral in coordinating arousal and can directly affect the activity of DA neurons^8,9^, how OX signaling influences DA release across states has not been investigated. The activity of OX neurons is lowest during NREM sleep, increased during REM sleep, and highest during wakefulness^50^. We predicted that OX neuron activity may recruit monoaminergic systems, including VTA DA, in the initiation or maintenance of both REM sleep and wakefulness. Surprisingly, in regions of the striatum, this is not the case, as loss of the OX system has no effect on DA release in the NAcc within states or across state transitions. However, it may be possible that during wakefulness, there are OX-dependent changes in DA release during behaviors such as feeding, drinking, or locomotion^1,51,52^. It may also be the case that while the loss of OX does not affect DA release in the striatum, it may affect DA release in other regions of the brain. We must also acknowledge that though DA release patterns were similar in both WT and OX KO mice, OX KO mice may have compensatory responses to the total, lifelong loss of OX signaling, and it is possible that more acute manipulations of the OX system could still modulate DA dynamics in the striatum across sleep-wake states. Nonetheless, our data suggest that alterations to DA release in the striatum are not responsible for EDS and DNS in narcolepsy.

Though we show that striatal DA release largely follows the activity of midbrain DA neurons, there are also notable differences. While VTA DA neuron activity increases prior to NREM-to-Wake transitions^16^, we show that across the DLS, NAcc, and NAcSh, increases in DA release only occur after the transition. This may indicate that DA release in the striatum contributes to the maintenance, but not generation, of wakefulness. The discrepancy in DA neuron activity and DA release may be accounted for by the fact that different populations of midbrain DA neurons and their respective projections may have different activity patterns across the sleep-wake cycle^18,53^. It is likely the case that distinct projections from midbrain DA neurons serve different functions across sleep-wake states and as such, the activity of these projections may not necessarily reflect the activity of the entire population. In addition, DA release in the striatum is influenced by a number of factors beyond VTA and SNc DA neuron activity, including expression of DAT^54^, glutamatergic transmission^55^, and the activity of striatal cholinergic interneurons^56,57^.

Previously, DA release has been evaluated in the NAc and DLS of WT mice. Hasegawa et al. demonstrated that, in the ventral NAc, analogous to our NAcSh, while robust dynamics were observed during REM sleep, optogenetic stimulation had no effect on REM sleep architecture^53^, which we also show to be true in OX KO mice. In the DLS, these studies concluded that DA release in the DLS was lowest during REM sleep relative to both wakefulness and NREM sleep^14^. We show here that, likely due to artifacts due to blood brain volume and pH^39^, this is not the case and that DA release in the DLS during REM sleep is elevated relative to NREM sleep. Furthermore, we show that while DA release is elevated across multiple striatal regions during REM sleep, only high frequency stimulation of the NAcc was sufficient to alter REM sleep latency, highlighting a specific role for the NAcc in the regulation of REM sleep. As such, further investigation is needed to determine the functional role of DA release in the NAcSh and DLS during REM sleep.

### Ventral striatal DA release increases prior to cataplexy

The limbic system has a documented role in regulating cataplexy^2,53,58–60^, however, the involvement of the striatum has garnered only modest attention^35,61^. In agreement with hypothesized roles for DA release in the striatum, we show that DA release in the DLS is suppressed during cataplexy, perhaps due to the suppression of movement. In the NAcc and NAcSh, robust increases in DA release were observed prior to cataplexy onset, consistent with the positively valenced states that often precede both human and murine cataplexy. Importantly, we show that this increase is a fundamental aspect of all cataplexy episodes, as the increase was independent of the behavior that preceded cataplexy. The relationship between DA release in the NAc and ‘cataplexy-like’ events has been previously explored in a non-narcolepsy mouse model (VTA terminal stimulation in the NAc using a step-function opsin in DAT-Cre mice), in which stimulation caused no change in the propensity of ‘cataplexy-like’ events^53^; this study then concluded that DA release in the NAc was not involved in regulation of cataplexy. However, in addition to not performing these experiments in a murine model of narcolepsy, optogenetic stimulation was primarily conducted in the NAcSh, which we also demonstrate shows no change in cataplexy frequency during stimulation in OX KO mice. We show that specifically stimulation of the VTA→NAcc circuit in OX KO mice is indeed sufficient to increase cataplexy propensity, highlighting a novel mechanism of cataplexy regulation.

As opto-/chemogenetic activation of the NAcSh is sufficient to increase cataplexy frequency in OX KO mice^35,61^, it remains unclear what mechanisms, if not DA, are responsible for this effect. While the NAcc and NAcSh share functional similarities, anatomically, the NAcSh has characteristics not present in the NAcc, such as a flipped patch-matrix, increased D1/D2-receptor co-expression, and differences in where DA axons synapse^27^. Additionally, as our viral injection strategy was not DA neuron-specific, activation of glutamatergic/GABAergic VTA projections may confound the ability to identify the specific role for DA. Indeed, the NAcSh in particular receives a high degree of glutamatergic input from the VTA^42,62^, while VTA GABA neurons directly project to cholinergic interneurons in the striatum, which themselves alter striatal medium spiny neuron activity^63,64^. Other monoaminergic systems (e.g., 5HT, NE)^65–67^ and limbic regions (e.g., CeA, mPFC, BLA) have also been implicated in cataplexy. As VTA→NAcc stimulation did not increase direct entrances into cataplexy, coincident recruitment or suppression of these systems may be required in addition to DA release in the NAcc to elicit cataplexy.

Midbrain DA neurons have typically been characterized to have two primary firing states: low-frequency tonic activity and high-frequency phasic activity, with these states thought to encode differing information^42^. Furthermore, sustained tonic DA release in the ventral striatum is thought to activate terminal D2 autoreceptors, thus suppressing phasic DA release^43–48^. We show that tonic stimulation of VTA efferents decreases phasic DA activity and suppresses both cataplexy and REM sleep. Phasic DA could contribute to cataplexy and REM sleep through activation of D1-expressing medium spiny neurons in the NAcc and NAcSh^15^, which directly project to^68^, and can inhibit, the ventrolateral periaqueductal gray and adjacent lateral pontine tegmentum (vlPAG-LPT), a region known to be REM-suppressive^69^. This would in turn provide disinhibition of the sublaterodorsal nucleus (SLD), a brainstem region integral in regulating muscle atonia during both REM sleep and cataplexy^50,70–73^. The OX system provides direct and indirect activation of the vlPAG-LPT^5^, but in the absence of a functional OX system, inappropriate inhibition of this circuit may underlie the sudden onset of muscle atonia in cataplexy (Figure 7G), as well as contribute to muscle atonia during REM sleep. As previously mentioned, other brain regions and neuromodulatory circuits are known to contribute to cataplexy and periods of intense emotion or positively valenced stimuli likely recruit these circuits, in addition to VTA DA neurons, to trigger cataplexy onset.

## Conclusion

Here, we show that the OX system regulates DA release in the limbic system but not during sleep-wake states. Moreover, we reveal that DA release in the ventral striatum is a key feature of both REM sleep and cataplexy, suggesting a link between DA release in the limbic system and muscle atonia-generating mechanisms. Together, these findings significantly improve our understanding of the neurobiology of narcolepsy. A more thorough understanding of the neural circuitry regulating reward processing, sleepiness, and muscle atonia will assist in developing novel, effective treatments of narcolepsy-cataplexy and other disorders resulting in EDS, DNS, and abnormal motor control across states, such as REM-sleep behavior disorder and obstructive sleep apnea.

## Methods

### Mice

Orexin knockout mice were generated from breeders obtained from Jackson Laboratory (B6.129S6-Hcrt*^tm1Ywa^*/J). Mice were genotyped using PCR with genomic primers 5’-GACGACGGCCTCAGACTTCTTGGG, 3’-TCACCCCCTTGGGATAGCCCTTCC, and 5’-CCGCTATCAGGACATAGCGTTGGC (with forward primers being specific for either wildtype or knockout mice and the reverse primer being common to both). All mice were housed in a University of Michigan vivarium in a temperature-controlled environment (12 h light and 12 h dark cycle; lights on at 2 AM) with *ad libitum* access to food and water. All animal protocols were approved by the University of Michigan’s Institutional Animal Care and Use Committee and are in accordance with NIH guidelines for the use and care of Laboratory mice. Mice that were used in the present study were maintained on a C57BL/6 background. Both male (WT, n = 3; OX KO, n = 12) and female (WT, n = 4; OX KO, n = 11) mice at least 10 weeks of age were used in all experiments. Overall health of experimental mice was monitored on a daily basis and any mice that displayed obvious signs of distress or weakness were removed from the study.

### Immunohistochemistry

Brain sections were washed in 0.1 M phosphate buffered saline pH 7.4, blocked in 3% normal donkey serum/0.25% Triton X100 in PBS for 1 h at room temperature and then incubated overnight at room temperature in blocking solution containing primary antiserum (rabbit anti-GFP, 488-conjugated, Novus #NB600-308, 1:5000; rabbit anti-TH, Thermo Fisher Scientific #OPA1-04050, 1:5000). The next morning, sections were extensively washed in PBS before being mounted onto polarized slides. For TH staining, sections were first washed in PBS/.05% Tween-20, then washed in PBS before being incubated in Alexa fluorophore secondary antibodies (donkey anti-rabbit, Thermo Fisher Scientific #A-21206, 1:1000) for 1 h at room temperature. After several washes in PBS, sections were mounted onto polarized slides and fluorescent images were captured with a Keyence BZ-X810 slide scanner microscope. All primary and secondary antibodies used are validated for species and application (1DegreeBio and Antibody Registry).

### Surgery

#### AAV injection

Mice were deeply anesthetized by inhalation of 2% isoflurane and placed on a Kopf stereotaxic apparatus (Tujunga, CA). Following standard disinfection procedure, the scalp was removed to expose the skull and a small hole was drilled into the skull unilaterally at defined positions to target NAcc (A/P: 1.2 mm, M/L: −1.3 mm, D/V: −4.1, −4.5 mm relative to bregma), NAcSh (A/P: 1.4: mm, M/L: −.7 mm, D/V: −4.6. −4.9 mm), DLS (A/P: 1 mm, M/L: −2.1 mm, D/V: −2.8, −3 mm), and VTA (A/P: −3.3, M/L: −.4, D/V: −4.3). A pulled-glass pipette with a ~20 µm tip diameter was inserted into the brain, and the virus was injected by an air pressure system. A picospritzer was used to control injection speed at 25 nl per min and the pipette was withdrawn 5 min after injection. For fiber photometry experiments, 300 uL of AAV5-CAG-dLight1.1 (AddGene #111067-AAV5; titer 7 × 10^12^ genome copies per mL) or AAV9-hSyn-eGFP (AddGene #50465-AAV9; titer 7 × 10^12^ genome copies per mL) was injected into the striatal region of interest (200 nL injected at the more ventral DV coordinate, 100 nL injected at the more dorsal DV coordinate). For optogenetic experiments, dLight1.1 was mixed with AAV6.2-CMV-Cre (AddGene #105537-AAV6.2; titer 1 × 10^13^ genome copies per mL) in a 4:1 mixture and injected into either the NAcc or NAcSh, and 200 uL AAV5-hSyn-Flex-ChrimsonR (AddGene #62723-AAV5; titer 5 × 10^12^ genome copies per mL) was injected into the VTA.

#### Optical fiber implantation

Optical fiber implantations were performed during the same surgery as viral injection (above). After viral injection, a metal ferrule optic fiber (400-µm diameter core; BFH37-400 Multimode; NA 0.37; ThorLabs) was implanted unilaterally over NAcc (A/P: 1.2 mm, M/L: −1.3 mm, D/V: −4.1 mm), NAcSh (A/P: 1.4 mm, M/L: −.7 mm, D/V: −4.6 mm), or DLS (A/P: 1 mm, M/L: −2.1 mm, D/V: −2.8 mm). Fibers were fixed to the skull using dental acrylic; after the completion of the experiments, mice were sacrificed, and the locations of optic fiber tips were identified based on the coordinates of Franklin and Paxinos^74^.

#### EEG/EMG implantation

EEG/EMG implantation was performed during the same surgery as viral injection and optical fiber placement (above). Mice were implanted with three stainless-steel screws (Frontal: A/P: 1.5 mm, M/L: −1.5 mm; Temporal: A/P: −3.5 mm, M/L: −2.8 mm; Ground: A/P: −3.3 mm, M/L: −.4 mm) and pair of multi strand stainless steel wires inserted into the neck extensor muscles. The EEG and EMG leads were wired to a small electrical connector that is attached to the skull with dental cement. Mice were then kept on a warming pad until awake and fed a regular chow (5L0D, LabDiet) diet throughout the experimental period unless otherwise noted. All mice were given analgesics (5 mg/kg Meloxicam) prior to the end of surgery and 24 hours after surgery. Mice were given a minimum of 2 weeks for recovery and 1 week for acclimation before being used in any experiments.

#### Polysomnographic recording and analysis

Polysomnographic signals were digitized at 1000 Hz, with a 0.3-100 Hz bandpass filter applied to the EEG and a 30-100 Hz bandpass filter applied to the EMG (ProAmp-8, CWE Inc.), with a National Instruments data acquisition card and collected using a custom MATLAB script. EEG/EMG signals were notch filtered at 60 Hz to account for electrical interference from the recording tether. Mice were recorded for a minimum of 3 sessions: a 12 h dark cycle recording, a 12 h light cycle recording, and a 12 h dark cycle where at the start of the recording, mice were given half of a Hershey’s™ kiss. Polysomnographic signals were analyzed using AccuSleep^75^, an open-source sleep scoring algorithm in MATLAB, and verified by an experienced sleep-scorer (BAT). Behavioral states were scored in 5 s epochs as either wake, NREM, REM, or cataplexy. Brief arousals were defined as transitions from NREM to wake that lasted <20 s. In subsequent photometry analysis, for primary state transitions, only transitions in which at least 30 s of a state occurred on either side of the transition were included. Cataplexy was defined by established guidelines and verified with videography^76^.

For assessment of behavior that occurred prior to the onset of cataplexy, overhead videography (9 fps) was acquired for each recording session. After initial scoring of “cataplexy-like” events using polysomnographic signals, videos were extracted around putative cataplexy events which contained the 40 seconds prior to the event, the entire cataplexy bout, and 40 seconds after the event. The behavior that occurred immediately prior to cataplexy was recorded, of which we determined six independent behaviors: eating, licking, running wheel, burrowing, grooming, and spontaneous (defined as the lack of a quantifiable behavior). Videos were scored by two independent observers (BAT, KSC). Unless otherwise noted, all cataplexy analysis was performed on data collected during the 12 h dark cycle in which chocolate was available, as the total amount of cataplectic events was highest during these recordings and there was no discernible difference in DA release dynamics when compared to recordings with no chocolate (Figure 2B-D). For both chocolate and no chocolate recordings, we required at least 5 episodes of cataplexy within the recording session to be included in the final group analysis. Recordings which did not meet this criterion were excluded from further analysis of dLight recordings during cataplexy but were still used for general sleep state analysis. When quantifying the average fluorescence within cataplexy episodes, we used the first 10 s of the episode.

The power spectral density of the EEG was computed using Welch’s method with a 1 s sliding Hanning window with 50% overlap and binned at 0.5 Hz resolution, from 0 - 15 Hz. In a subset of mice, EEG artifacts confounded calculation of the power spectrum. To address this, within each bout of a respective state, the average power was calculated, and outlier detection was performed using the MATLAB function *rmoutliers*. Briefly, bouts were excluded from the final average if the mean power of that bout was greater than three scaled median absolute deviations from the median. To facilitate comparisons across mice, spectra were normalized: (*norm(x) = (x − min(x)) / (max(x) − min(x))*). In Figures 3 and 5, theta power was calculated using the filter-Hilbert method. EEG signals were first filtered at the theta range (5 - 9 Hz) and separately between 0 - 30 Hz. The Hilbert transform was applied to the absolute value of these filtered signals, after which the theta signal was normalized by dividing this signal by the 0 - 30 Hz signal.

### *In vivo* fiber photometry

#### Acquisition and post-processing

Beginning 2-3 weeks after surgery, mice were connected to an EEG/EMG recording tether and a fiber optic patch cable and given 5-7 days to habituate to the recording environment, which includes a food hopper (FED3)^77^, capacitive lickspout, and IR-based running wheel for automated behavioral readouts. Mice were considered fully habituated after having built a nest and were comfortably using the in-cage FED3, lickspout, and running wheel. Both the lickspout and running wheel were designed using custom Arduino code. Mice were able to move freely within the behavior chamber while outfitted with both optic fibers and EEG/EMG tethers. Fiber optic patch cables (0.8 - 1 m long, 400 μm diameter; Doric Lenses) were firmly attached to the implanted fiber optic cannulae with zirconia sleeves (Doric Lenses). LEDs (Plexon; 473 nm) were set such that a light intensity of <0.1mW entered the brain; light intensity was kept constant across sessions for each mouse. Emission light was passed through a filter cube (Doric) before being focused onto a sensitive photodetector (2151, Newport). Signals were digitized at 60 Hz using PyPhotometry^78^, which allows for pulsed delivery of light, minimizing the amount of bleaching over the course of 12 hr recordings.

To address photobleaching over the course of the recording period, the photometry signal was corrected by subtracting a double exponential fit, then adding back the mean of the trace. Signals were then smoothed with a 120 ms sliding window and background autofluorescence was subtracted. For each recording session, the smoothed, detrended photometry signal was converted to ΔF/F ((F – F_0_)/F_0_); where F_0_ was calculated as the 10th percentile of the entire fluorescence trace). These traces were then z-scored using the MATLAB *zscore* function to facilitate comparisons across days and mice. Slow drift was removed from the z-score-transformed ΔF/F values using the MATLAB script ‘BEADS’ (as used by others^79,80^) with a cutoff frequency of either 0.00035 cycles per sample (for transition analysis) or 0.0035 cycles per sample (for transient analysis), filter order of 2, and an asymmetry parameter of 5.

#### Analysis within sleep states

We detected transients using a method similar to that reported in previous studies^16,81^. Briefly, we generated two filtered ΔF/F signals: one low-pass filtered at 0-4 Hz, and the other low-pass filtered at 0-40 Hz. The derivative of their squared difference was calculated, and candidate transients were detected by thresholding this signal at the mean + 2 s.d. Times of the candidate transients were extracted and compared with the original ΔF/F: transients in which the ΔF/F at this time was greater than the mean + 2 s.d. were included in the final analysis. Lastly, the exact peak time of each transient was identified by taking the maximal value of the thresholded signal within each transient. For each session and for each sleep-wake state, we calculated the mean transient rate within each animal. For quantifying DA dynamics across state transitions, we identified all points of state transition and aligned the ΔF/F trace around these times. Only transitions in which there were at least 30 s of a state on either side of the transition were included in subsequent analysis.

Prior research has suggested that DA release in the NAcc should increase at the onset of REM sleep^15,18^. Yet when recording dLight fluorescence during REM, we observed a consistent decrease over the course of the bout, in spite of increased phasic DA release (Figure 1C-E; Figure S1I, J). To verify whether this was an artifact of fluorescent recording during REM sleep, we evaluated fluorescence in GFP-expressing mice as well. We found a similar reduction in fluorescence during REM, however, when comparing the GFP signal to the dLight signal, the GFP signal had a lack of phasic activity (Figure 1F-H; Figure S1M, N). As such, it is likely that the observed drop in baseline fluorescence during REM sleep is artificial and that analysis of transient activity is a better representation of ongoing DA activity during REM sleep. Additionally, during REM sleep, physiological changes in body temperature^82^, blood flow^83,84^, and pH^85^ occur, all of which may contribute to altering the fluorescent protein and could account for artifacts in the fluorescent signal. Recent publications have also demonstrated that changes in blood brain volume^39^ and hemoglobin absorption^38^ contribute to artifacts present in fiber photometry recordings, particularly during REM sleep. As such, specifically for analysis of DA release within REM sleep, we employed methods such as BEADS (see above) to remove this artifact, resulting in a more accurate representation of within state activity.

### *In vivo* optogenetic studies

#### Stimulation parameters

Following recording mice across the light and dark cycle with and without chocolate (as described above), mice expressing ChrimsonR underwent additional recording sessions, as follows. *5 s 20 Hz stimulation:* NAcc and NAcSh mice were stimulated with red light (625 nm) every 8-10 min at 20 Hz, 5 ms pulses for 5 s during the 12 h dark cycle (Figure S6A). *20 s 10-40 Hz variable stimulation:* NAcc and NAcSh mice were stimulated with red light with a variable frequency stimulation protocol. Stimulation began at 10 Hz, 5 ms pulses for 1 s, after which every s, the frequency increased by 2 Hz. This continued until 40 Hz, at which this frequency was held for 5 s (Figure S6A). Stimulation occurred with a variable 8-10 min ITI; this paradigm was intended to mirror the spontaneous dynamics of dopamine release during cataplexy and REM sleep. *2 Hz continuous stimulation:* NAcc and NAcSh mice were stimulated for 2 h between ZT15-19 at 2 Hz, 15 ms duration. This stimulation was intended to elevate tonic, rather than phasic, DA levels. Photostimulation was provided using a waveform generator (Arduino electronics platform) that provided TTL input to a red light laser (Thor labs). We adjusted the power of the laser such that the light power exiting the fiber optic cable was at least 10 mW/mm^2^.

#### Analysis of optogenetic stimulation

For phasic stimulation recordings, individual stimulation pulse times were acquired to facilitate analysis time locked to the beginning of stimulation trains. When calculating latencies to wake and REM, only stimulation that occurred after at least 30 s of stable (no interruptions from brief arousals) NREM were included in analysis. The same criterion applied to latency to cataplexy when stimulation occurred during wake. In these recordings, the “Stim” group was the group in which stimulation was applied throughout the 12 hr recording period. The “No stim” group was defined by overlaying the TTLs acquired during the stimulation recording to a 12 hr recording period done during the dark cycle with no chocolate or stimulation present for the same animal. For tonic stimulation recordings, all recordings were done between 15:00 - 19:00. For tonic stimulation, we required that the “No stim” recording had at least one episode of REM sleep or cataplexy to be included in the final analysis.

A possible confound of combining fiber photometry with optogenetics is that there is a small section of overlap between the excitation wavelength of ChrimsonR and the 473 nm light used to facilitate dLight recordings. As a result, it is possible that during fiber photometry recordings there is sustained low-level activation of ChrimsonR-expressing fibers in the striatum, which may induce behavioral changes. However, the observed overlap is minimal, with 473 nm light only accounting for ~25% of ChrimsonR’s maximal excitation^86^. Additionally, for fiber photometry, we adjust the laser power to be two orders of magnitude lower than that used for optogenetics, further reducing the likelihood of producing unwanted behavioral effects. As such, we do not expect that there were any significant behavioral changes in mice expressing both dLight1.1 and ChrimsonR in striatal regions.

### Statistical analysis

Statistical analyses were performed using Prism 9.0 (GraphPad) software. Details of statistical tests employed can be found in the relevant figure legends and an alpha level of 0.05 was used for determining significance. The data presented met the assumptions of the statistical test employed. Exclusion criteria for experimental mice were (i) sickness or death during the testing period (ii) if histological validation of the injection site demonstrated an absence of reporter gene expression (iii) or if histological validation demonstrated a mistargeted fiber placement. These criteria were established before data analysis. N numbers represent final numbers of healthy/validated mice.

## Acknowledgments

We thank Chris Phillips for initial assistance with fiber photometry, Timothy Cha for assistance with husbandry, and Dr. Liam Potter for helpful discussion. This work was supported by a University of Michigan Rackham Graduate Student Research Grant and Rackham Merit Fellowship (BAT), a Gilmore Award from the Gilmore Fund for Sleep Research and Education, an E. Matilda Ziegler Award, a Whitehall Foundation new investigator grant, and an NIH R01DK129366 (CRB).

## Author contributions

BAT contributed to the acquisition of data. BAT, KSC, and CRB contributed to the analysis and interpretation of the data. BAT and CRB contributed to drafting the manuscript.

