## Supplemental Figures for "Striatal dopamine regulates sleep states and narcolepsy-cataplexy"

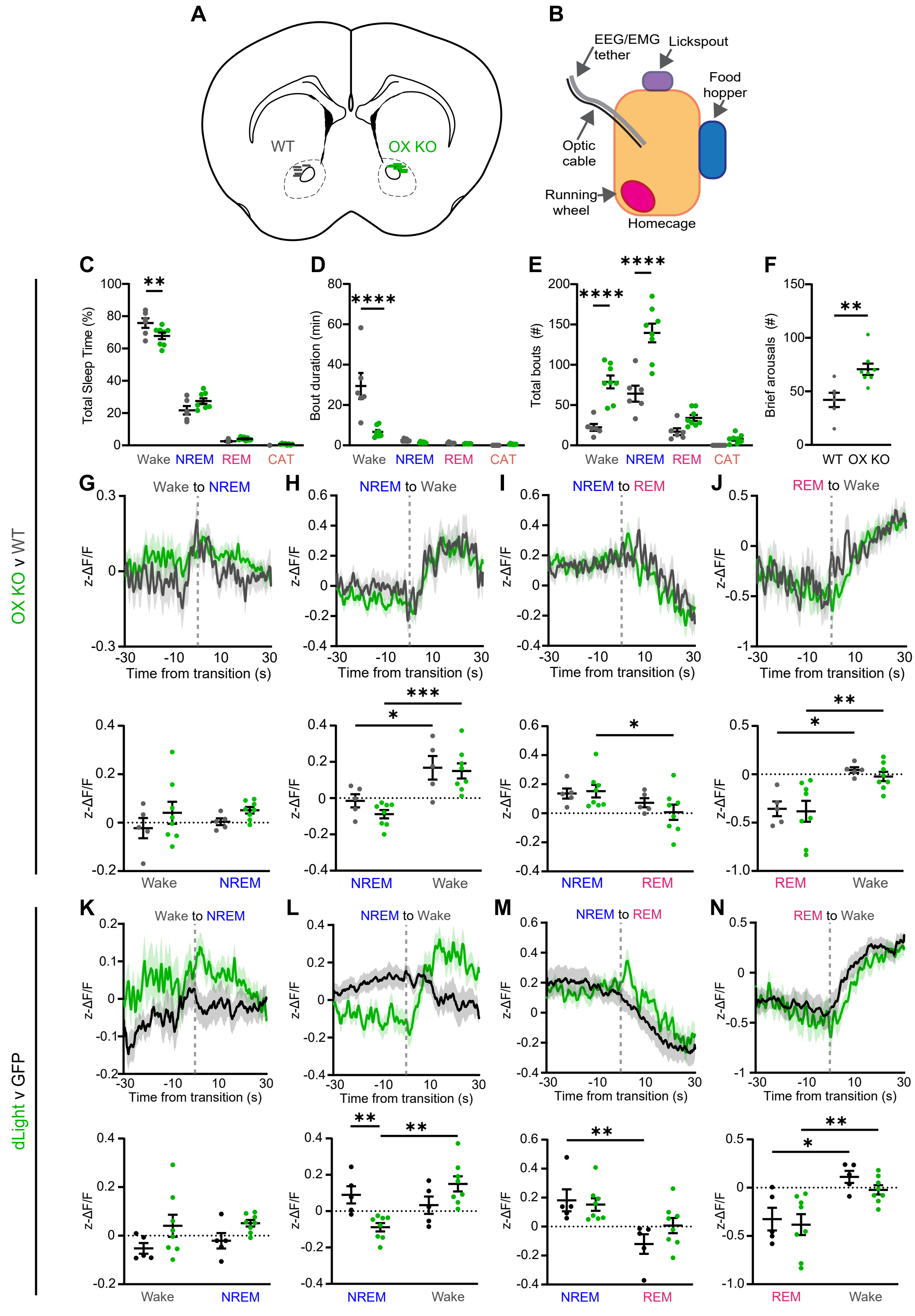


Figure S1. Related to Figure 1.

**Supplemental Figure 1. DA release in the NAcc fluctuates across sleep-wake transitions independent of OX expression. Related to Figure 1.**

(A) Optic fiber locations in the NAcc for WT (n = 5) and OX KO (n = 8) mice.

(B) Schematic representation of recording chambers outfitted with food hopper, lickspout, and running wheel. (C - E) Percentage of time spent (C), average bout duration (D), and total number of bouts (E) in wake, NREM, REM, and cataplexy during a 12 h dark cycle recording (WT, n = 6; OX KO, n = 8).

(F) Total number of brief awakenings from NREM.

(G - J) dLight fluorescence aligned to Wake-to-NREM (G), NREM-to-Wake (H), NREM-to-REM (I), and REM-to-Wake (J) transitions during a 12 h light cycle for WT (n = 5) and OX KO (n = 8) mice (top). Average dLight fluorescence of the 30 s before and after the transition (bottom).

(K - N) Average dLight fluorescence aligned to NREM-to-REM (K), REM-to-Wake (L), Wake-to-NREM (M), and NREM-to-Wake (N) sleep transitions during a 12 h light cycle for GFP- and dLight-expressing OX KO mice (top). Average dLight fluorescence of the 30 s before and after the transition (bottom).

All data represented as the mean ± SEM. *$p<0.05$, **$p<0.01$, ***$p<0.001$, ****$p<0.0001$; two-way ANOVA with Sidak post-hoc comparison test (C - E, G - N), unpaired t-test (F).

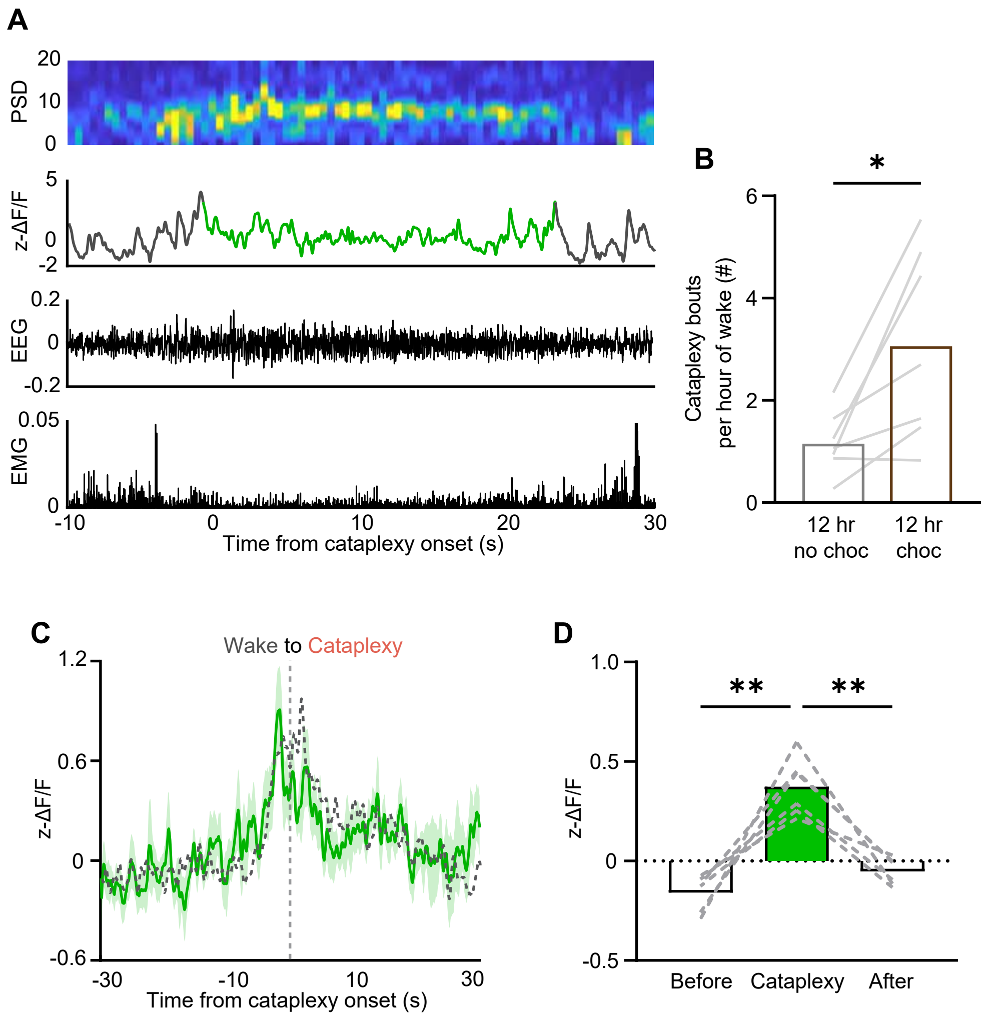


Figure S2. Related to Figure 2.

**Supplemental Figure 2. NAcc DA response during cataplexy does not change in the presence or absence of chocolate. Related to Figure 2.**

(A) Representative power spectrogram, ΔF/F trace, EEG, and EMG during a single cataplexy episode in a NAcc dLight-expressing OX KO animal.

(B) Comparison of average number of cataplexy bouts per hour of wakefulness between a 12 h dark cycle with and without chocolate availability (n = 7).

(C) Average dLight fluorescence aligned to Wake-to-Cataplexy transitions that occurred in the dark cycle without chocolate availability (solid green trace; dotted black traced overlayed represents the same transition that occurred with chocolate availability).

(D) dLight fluorescence in the period prior to, during, and after cataplexy that occurred in the absence of chocolate (n = 6).

All data represented as the mean ± SEM. *p<0.05, **p<0.01, ***p<0.001, ****p<0.0001; RM one-way ANOVA with Tukey post-hoc comparison test (D), paired t-test (B).

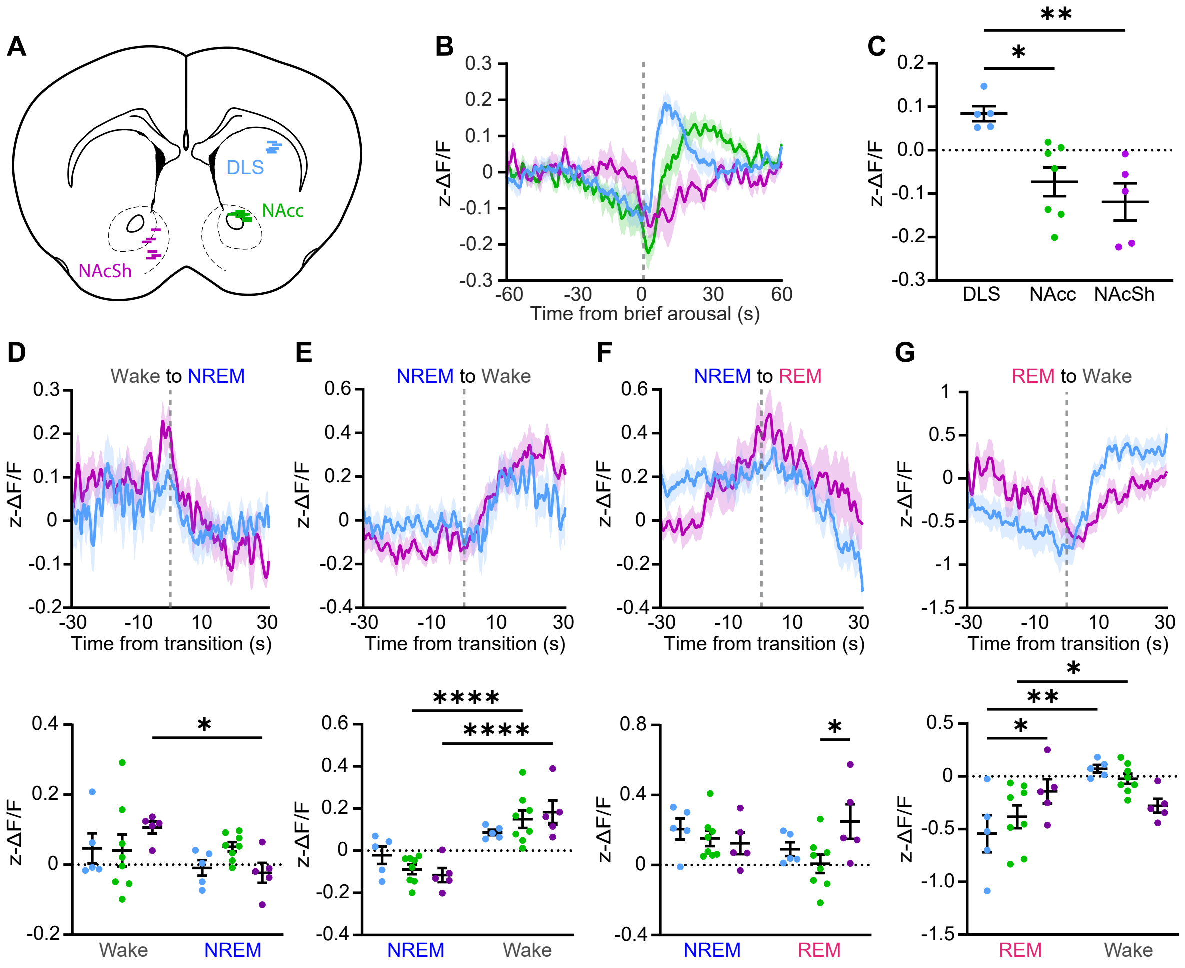


Figure S3. Related to Figure 3.

**Supplemental Figure 3. DA release differs across striatal regions during sleep-wake transitions. Related to Figure 3.**

(A) Location of optic fibers tips in relevant striatal regions.

(B) dLight fluorescence aligned to brief arousals from NREM.

(C) Average dLight fluorescence during brief arousals.

(D - G) dLight fluorescence aligned to Wake-to-NREM (D), NREM-to-Wake (E), NREM-to-REM (F), and REM-to-Wake (G) transitions during a 12 h light cycle (top).

Average dLight fluorescence of the 30 s before and after the transition (bottom). All data represented as the mean ± SEM. *$p<0.05$, **$p<0.01$, ****$p<0.0001$; one-way ANOVA with Tukey post-hoc comparison test (C), two-way ANOVA with Sidak post-hoc comparison test (D - E).

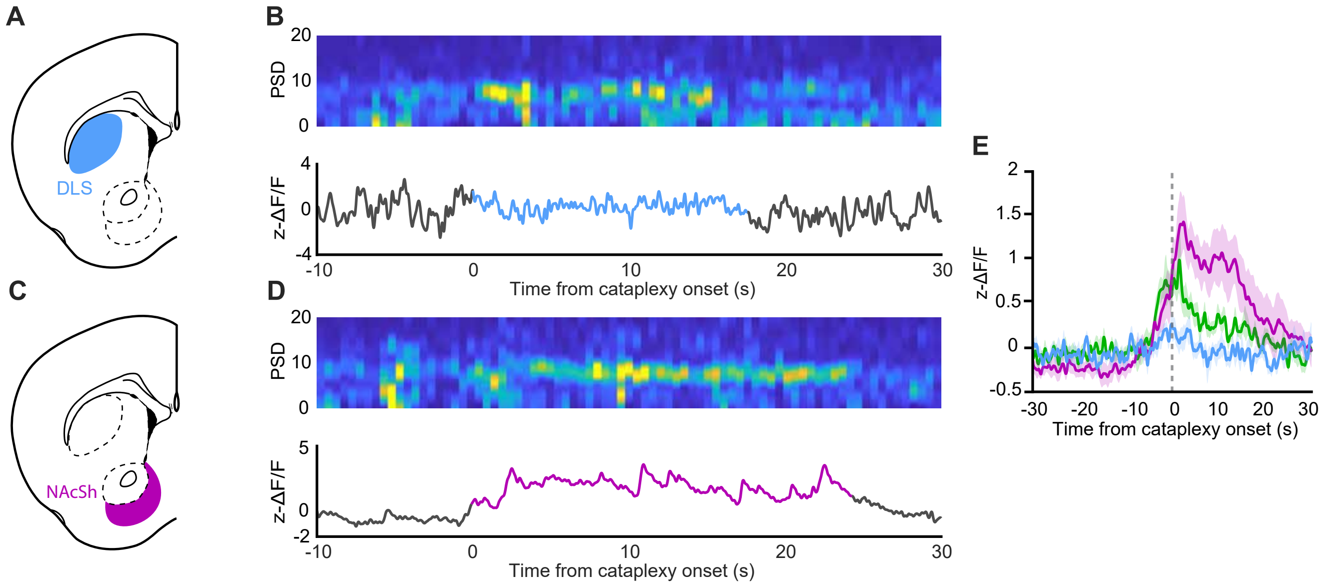


Figure S4. Related to Figure 4.

**Supplemental Figure 4. DA release in the NAcSh, but not the DLS, is elevated during cataplexy.**

(A) Schematic representation of the DLS.

(B) Representative power spectrogram and ΔF/F trace during a single episode of cataplexy in a DLS dLight-expressing OX KO animal.

(C) Schematic representation of the NAcSh.

(D) Representative power spectrogram and ΔF/F trace during a single episode of cataplexy in a NAcSh dLight-expressing OX KO animal.

(F) Overlay of average dLight fluorescence across striatal regions during Wake-to-Cataplexy transitions.

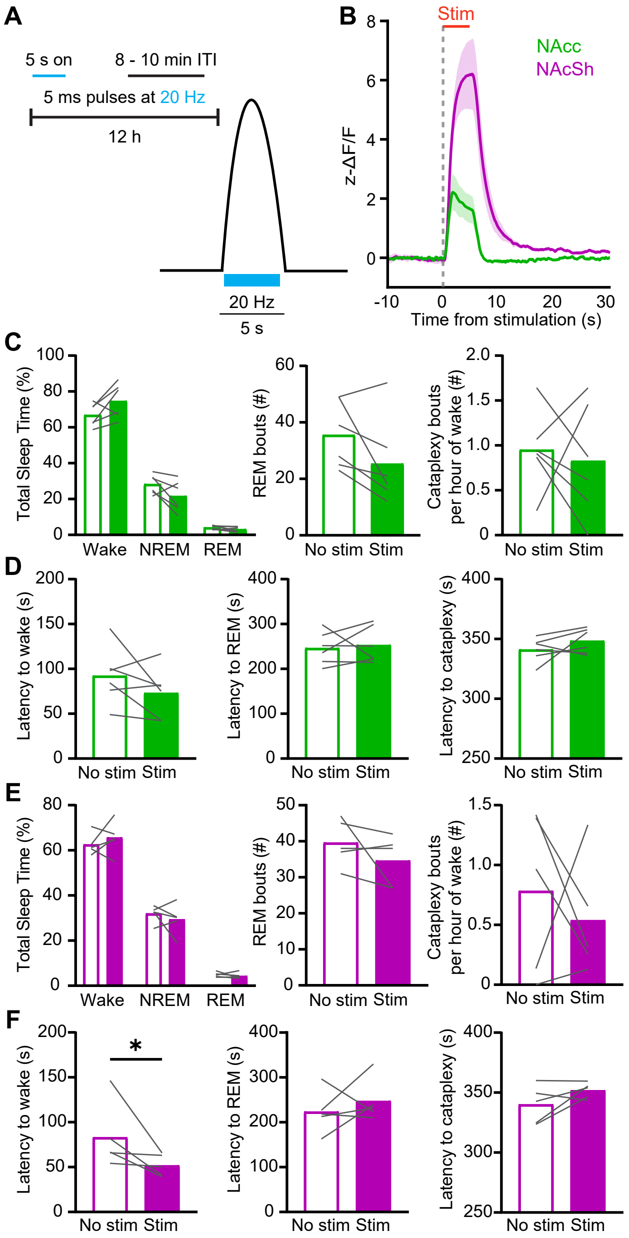


Figure S5. Related to Figure 7.

**Supplemental Figure 5. Brief phasic stimulation of VTA efferents in the NAcc and NAcSh has little effect on sleep and cataplexy. Related to Figure 7.**

(A) Schematic of the 12 h 20 Hz stimulation paradigm.

(B) dLight response to 20 Hz stimulation in the NAcc (n = 6) and NAcSh (n = 5).

(C) Effect of 20 Hz stimulation in the NAcc on total sleep time, number of REM bouts, and number of cataplexy bouts per hour of time spent awake.

(D) Effect of 20 Hz stimulation in the NAcc on latency to Wake, REM, and Cataplexy.

(E, F) Same as (C, D), but for the NAcSh.

(G - J) Effect of 20 Hz (G, I) and variable sweep (H, J) stimulation in the NAcc and NAcSh on the probability of Wake, NREM, and REM when stimulated during NREM.

All data represented as the mean ± SEM. $^*p<0.05$; paired t-test.

| | Wake | NREM | REM | References |
|---|---|---|---|---|
| **DLS** | DA↑↑<br>DA↑↑ | DA↓<br>DA↓ | DA↑<br>DA↓ | Toth et al., 2023<br>Dong et al., 2019 |
| **NAcc/NAcSh** | DA↑<br>DA↑<br>DA↑ | DA↓<br>DA↓<br>DA↓ | DA↑↑<br>DA↑<br>DA↑↑ | Toth et al., 2023<br>Lena et al., 2005<br>Hasegawa et al., 2022 |
| **VTA cell bodies** | Active: ↑↑ Quiet: ↑<br>Active: ↑↑ Quiet: ↑<br>Active: ↑↑ Quiet: ↑ | ↓<br>↓<br>↓ | ↓<br>↑↑<br>↑↑ | Trulson and Preussler, 1983<br>Dahan et al., 2007<br>Eban-Rothschild et al., 2016 |

**Supplemental Table 1. Summary of DA release in striatal regions across sleep-wake states.**

| | Stimulation | Inhibition | References |
|---|---|---|---|
| **NAcc** | Phasic: ↑Wake, ↑REM, ↑Cataplexy<br>Tonic: ↓REM, ↓Cataplexy<br>↑Wake | -<br>- | Toth et al., 2023<br>Toth et al., 2023<br>Eban-Rothschild et al., 2016 |
| **NAcSh** | No effect<br>No effect | -<br>- | Toth et al., 2023<br>Hasegawa et al., 2022 |
| **VTA cell bodies** | ↑Wake | ↑NREM | Eban-Rothschild et al., 2016 |

**Supplemental Table 2. Summary of the sleep effects of DA terminal stimulation/inhibition in striatal regions.**
